# Dynamic aspects of plant water potential revealed by a microtensiometer

**DOI:** 10.1101/2021.06.23.449675

**Authors:** Vinay Pagay

## Abstract

Water potential is a fundamental thermodynamic parameter that describes the activity of water. In this paper, we describe the continuous measurement of plant water potential, a reliable indicator of its water status, using a novel *in situ* sensor known as a ‘microtensiometer’ in mature grapevines under field conditions. The microtensiometer operates on the principle of equilibration of water potentials of internal liquid water with an external vapour or liquid phase. We characterised the seasonal and diurnal dynamics of trunk water potentials (Ψ_trunk_) obtained from microtensiometers installed in two grapevine cultivars, Shiraz and Cabernet Sauvignon, and compared these values to pressure chamber-derived stem (Ψ_stem_) and leaf (Ψ_leaf_) water potentials as well as leaf stomatal conductance. Diurnal patterns of Ψ_trunk_ matched those of Ψ_stem_ and Ψ_leaf_ under low vapour pressure deficit (VPD) conditions, but diverged under high VPD conditions. The highest diurnal values of Ψ_trunk_ were observed shortly after dawn, while the lowest values were typically observed in the late afternoon. Differential responses of Ψ_trunk_ to VPD were observed between cultivars, with Shiraz more sensitive than Cabernet to increasing VPD over long time scales, and both cultivars had a stronger VPD response than soil moisture response. On a diurnal basis, however, time cross correlation analysis revealed that Shiraz Ψ_trunk_ lagged Cabernet Ψ_trunk_ in response to changing VPD. Microtensiometers were shown to operate reliably under field conditions over several months. To be useful for irrigation scheduling of woody crops, new thresholds of Ψ_trunk_ need to be developed.

## Introduction

Measurements of crop water status, which are essential for optimised irrigation scheduling, have historically relied on low throughput and high cost instruments, and labour intensive methods, that have decreased their utility and uptake by the farming community. One such method widely established to reliably quantify plant water status is the manual measurement of leaf or stem water potential (Shackel, 2011; Williams and Baeza, 2007), using a Scholander pressure chamber invented in the 1960s (Scholander *et al.*, 1965). To overcome some of the limitations of such manual techniques, several electronic plant-based sensors to continuously measure crop water status have recently been developed that are based on various sensing modalities. These include sap flow sensors (Ginestar *et al.*, 1998), thermal diffusivity sensors (Pagay and Skinner, 2018), dendrometers (Corell *et al.*, 2014), and thermal or infrared sensors (Jones, 1999). A recent review of several of these sensors as applied in tree fruit crops can be found in Scalisi *et al.* (2017).

Given the prevalence of water potential as a reliable crop water status metric, sensors to measure plant water potential have also been developed. These sensors, known as hygrometers or psychrometers, provide continuous measurements of water potential either as contact sensors on leaves or *in situ* sensors embedded in stems (Dixon and Tyree, 1984; McBurney and Costigan, 1984; Michel, 1977). Hygrometers measure the water potential of the vapour phase. They are prone to significant errors due to the requirement of isothermal conditions between the measurement junction and plant tissue (Dixon and Tyree, 1984). A 1 °K temperature difference between the plant tissue and sensor can result in a water potential error of over 7.7 MPa (Dixon and Tyree, 1984). Much like stem hygrometers, other *in situ* sensors embedded in the stems or trunks include those that measure the osmotic potential of the xylem tissue and calibrated to stem water potential (Meron *et al.*, 2015). The osmotic potential sensor requires proper fluidic contact between the sensor and the plant tissue, as well has long transients (order hours) associated with the measurement.

Tensiometers measure the water potential of an external matrix by equilibrating an internal, constant volume of water whose hydrostatic pressure is taken as the negative of the external water potential. Tensiometers were originally developed for measurement of soil matric potential (Richards, 1942) and have been used for irrigation scheduling of crops (Cormier *et al.*, 2020). Based on this principle, Pagay *et al.* (2014) developed MEMS-based tensiometers, so-called ‘microtensiometers’ (MT), for rapid measurements of the water potential of an external matrix. The MTs were previously shown to operate reliably down to below −10 MPa with short transients (equilibration or response times) of ~ 20 min. This measurement range and temporal resolution makes the sensors valuable for not only crop and soil water status monitoring, but also other contexts including meteorology, concrete curing and food processing, and other systems where the internal water status is required. Subsequent improvements on the original MT design for improved transients (faster response times) were made by Black *et al.* (2020). This second generation microtensiometer was used for both *in situ* and *ex situ* measurements of water potential in a range of matrices including foods.

This paper presents the first results of field experiments with MTs, embedded water potential sensors, in mature grapevines grown under different environmental and soil moisture conditions in a Mediterranean climate. We compared the dynamic MT responses of plant water potential to values of leaf and stem water potentials as measured by the Scholander pressure chamber over both long term and short term (diurnal) periods, and with other environmental metrics including soil moisture and atmospheric conditions. Our goal was to validate the use of MTs in a field context and to provide new insights into the dynamics of plant water potential.

## Materials and Methods

### Experimental site and plant material

Two commercial vineyard blocks located in the Coonawarra region of South Australia (37.29° S, 140.83° E) were selected for the trial. One block was planted in 1988 to *Vitis vinifera* cv. Cabernet Sauvignon grafted onto Schwarzmann rootstock, while the second block was planted in 2013 to *V. vinifera* cv. Shiraz (syn. Syrah) grafted onto Teleki 5C rootstock. Both vineyards were situated within 5 km of each other and planted over the dominant ‘terrarossa’ soil, characterised by a distinctive red-brown, thick, clay B horizon soils overlying limestone, the depth to which is variable. In each vineyard, three adjacent vines per cultivar were selected for measurements, and, additionally, the middle vine for continuous monitoring of soil and plant water status (see details below). Vineyard management, irrigation, and integrated pest management were applied to both blocks as per convention in the region for premium winegrape production.

### Environmental monitoring and climatic conditions

Environmental (weather) data for the vineyard blocks was obtained from the Coonawarra weather station maintained by the Australian Bureau of Meteorology (BOM), Station ID: 026091. Daily maximum air vapour pressure deficit (VPD) was calculated using maximum temperature and minimum relative humidity (RH) daily data. The long-term (20-year) mean January temperature (MJT) for Coonawarra is 19.3°C and the growing degree days (GDD; base 10 °C; October-April) is 1511. The climate of the area is characterised as Mediterranean, with winter dominant rainfall and relative summer drought. Average annual rainfall for Coonawarra is approx. 569 mm (Bureau_of_Meteorology, 2021). Supplemental irrigation is typically required from December until March. The elevation of the region is between 57 m and 63 m above sea level.

### Soil moisture measurements

Of the three sentinel adjacent vines in each cultivar/block, the middle vine was selected for continuous monitoring of soil moisture, temperature and electrical conductivity using a capacitance-based sensor (Model: Teros-12, Meter Group, Pullman, WA, USA) buried approx. 30 cm below the surface and approx. 10 cm from the trunk of the vine in the vine row. The hourly sensor data was wirelessly transmitted via telemetry to a Cloud-based server and visually displayed on a user interface (Greenbrain, Measurement Engineering Australia, Adelaide, SA, Australia).

### Plant water status measurements

#### Leaf stomatal conductance

In each block, leaf stomatal conductance (*g*_s_) was measured on the three sentinel vines per cultivar between 1200 – 1300 h. Measurements were performed on one fully-expanded, healthy leaf per vine using an open system infrared gas analyser (IRGA; LI-6400XT, LI-COR Biosciences Inc., Lincoln, NE, USA) with a 6 cm^2^ cuvette. An external LED light source (LI-6400-02B) attached to the cuvette was used at a fixed PAR value of 1500 μmol m^−2^ s^−1^ due to the sometimes variable ambient light levels. The cuvette gas flow rate was set at 400 μmol s^−1^ and reference CO_2_ was set to 400 ppm. The cuvette and leaf temperatures were at ambient (uncontrolled), while cuvette relative humidity with the leaf inserted was maintained within a range of 35-55%. After IRGA measurements were performed, the same leaf was excised to determine leaf water potential (see below). IRGA measurements were conducted diurnally (every 2 h between 0800 and 2000 h) on two days of contrasting VPDs – high VPD (February 17, 2021) and low VPD (January 26, 2021) – that were typical of a Mediterranean region.

#### Leaf and stem water potentials

In each block, midday stem (Ψ_stem_) and leaf water potentials (Ψ_leaf_) were measured on adjacent mature leaves of the same shoot between the hours of 1200-1300 using a Scholander pressure chamber (Soil Moisture Equipment Corp., Santa Barbera, CA, USA). For Ψ_stem_ measurements, leaves were bagged using an opaque aluminium-lined bag for a minimum of one hour prior to measurement to stop transpiration and allow for equilibration of water potentials between the leaf and shoot (stem). Measures of Ψ_leaf_ and Ψ_stem_ were performed on one leaf per vine from the three sentinel vines in each block/cultivar. Leaf and stem water potentials were measured diurnally on the two measurement days concurrently with IRGA measurements. The same leaf used for *g_s_* measurement with the IRGA was used to measure Ψ_leaf_.

#### Trunk water potentials

On December 12, 2020, a total of four microtensiometers (MT; FloraPulse, Davis, CA, USA) were embedded into the trunks of the grapevines, two MTs per vine per cultivar. Readers are referred to Pagay *et al.* (2014) for a detailed description of the theory of tensiometry, and to Black *et al.* (2020) for technical details of the MT sensor design, fabrication, calibration, and lab testing results (performance under controlled conditions). Briefly, the MT consists of a micromachined piezoelectric pressure sensor coupled to a nanoporous silicon membrane via a cavity of liquid water. The membrane couples the external environment with the internal sensor water, allowing for the equilibration of water potentials. Decreases in the external water potential, i.e. below saturation vapour pressure, results in decreases in the hydrostatic pressure of the internal water (*P_liq_*), as the liquid phase water comes under tension or is stretched (*P_liq_* < 0.1 MPa). This relationship is shown in Eqn. 1 below.

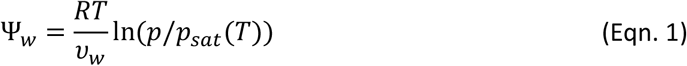

where, Ψ_w_ is the water potential (= *P_liq_* – *P_atm_*), *R* is the ideal gas constant (8.314 J mol^−1^ K^−1^), *v_w_* is the molar volume of water (1.8 x 10^−5^ m^3^ mol^−1^), *p* is the partial pressure of vapour, and *p_sat_* is the saturation vapour pressure at temperature, *T*. The ratio of *p/p_sat_* is the relative humidity of the sample.

Two MTs (Fig. 1a) were embedded into each trunk of a woody, mature grapevine per cultivar. Sensor installation consisted of the following steps: (1) removal of bark on a flat section of the trunk; (2) removal of phloem tissue using a cork borer and spatula/blade; (3) insertion of a custom stainless steel sleeve (Fig. 1b; OD: 14 mm, ID: 9 mm) using a hammer; (4) Drilling into the sleeve approx. 5 cm into the trunk (within xylem tissue) and removing tissue; (5) filling the cavity with a kaolin-based mating compound; (6) inserting the hydrated MT into the mating compound; (7) placing a stainless steel cap to close the sleeve; (8) Covering the sleeve exterior at the trunk with silicone to ensure air and water proofing; (9) wrapping plastic film around the sensor followed by a 25 mm thick foam batting with reflective aluminium film to minimise exterior temperature fluctuations to the sensors (Fig. 1c). The sensors equilibrated with the vine (through the mating compound) within two days of installation. The MT data of trunk water potential (Ψ_trunk_; value averaged for both sensors), was obtained every 20 min wirelessly transmitted via telemetry to a Cloud-based server (Amazon Web Services, USA) and visually displayed on a user interface (FloraPulse, Davis, CA, USA).

**Figure 1:**
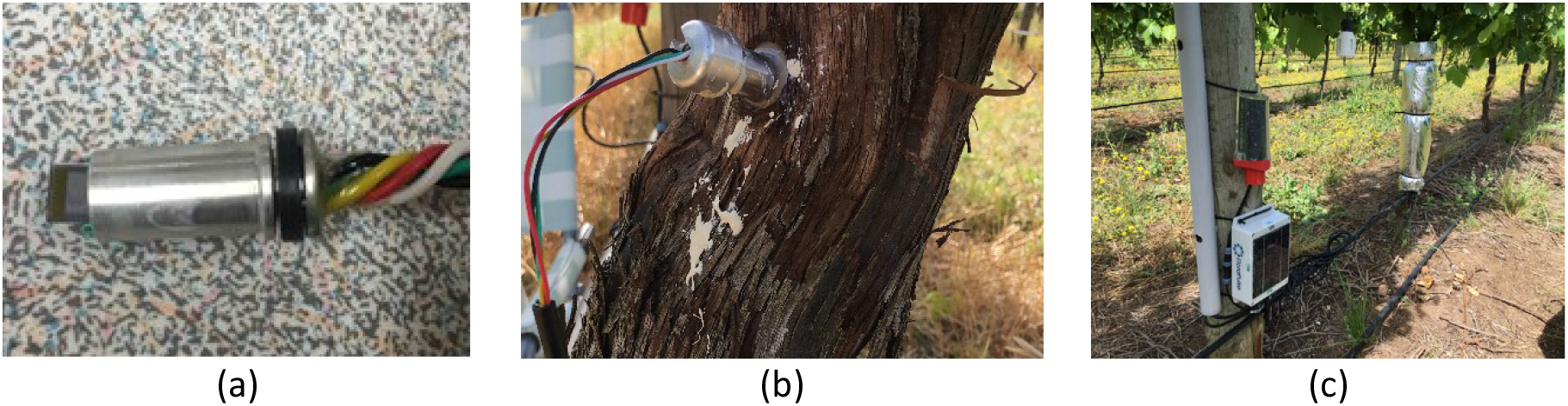
**(a)** Close up view of an individual microtensiometer showing the stainless steel packaging surrounding a protruding sensor chip. The tip of the chip consists of the nanoporous silicon membrane that allows for equilibration of water potentials between the exterior matrix and internal water; **(b)** MTs installed in a grapevine trunk within a stainless steel sleeve filled with kaolin mating compound; **(c)** Background: MT batting and reflective film to minimise temperature effects on water potential measurements of the sensor; Foreground: dataloggers and wireless transmitters of the MT (bottom unit, white box) and soil moisture sensor (top unit, red base).

### Statistical analyses

Time-lagged cross correlation (TLCC) analysis was used in MATLAB programing software (v.9.8.0, R2020a, The MathWorks, Inc., Natick, MA, USA) to analyse the continuous (20-min interval) data of VPD and Ψ_trunk_ over the course of two days with contrasting VPDs: January 26, 2021 (low VPD day; daily max. VPD ~ 1.6 kPa) and January 24, 2021 (high VPD day; daily max. VPD ~ 6.7 kPa). TLCC analysis involves determining the correlations between two time series datasets that are shifted in time (Chatfield and Xing, 2019), and repeatedly calculating Pearson Product Moment Correlation (cross correlation) Coefficient (XCC) after each shift (Cheong, 2020). Resulting ‘offset’ values, which when selected at the highest normalised XCC in the series, indicate the time shift (lag or advancement) of a particular time series compared to the other. Time shifts were selected such that they aligned with the VPD and Ψ_trunk_ measurement interval of 20 min, therefore each offset represented 20 min.

## Results

### Environmental conditions and soil moisture

Measurements of environmental conditions and vine water status were made over the course of two days, one with high vapour pressure deficit (VPD) and the other with low VPD, during the 2020-21 growing season from 0800-2000 h. The low VPD day, January 26, 2021, was characterised by sunny and cool conditions, with the daily maximum temperature reaching just over 23 °C with a maximum VPD of 1.7 kPa at around 1400 h (Fig. 2 a,g). This VPD was maintained until around 1600 h after which a decrease in solar radiation and temperature resulted in a decrease. Photosynthetically active radiation (PAR) was highest earlier in the day, around 1100 h, and declining gradually after this time. On the high VPD day, February 17, 2021, maximum daily temperature was nearly 37 °C and maximum VPD was approx. 6.0 kPa (Fig. 2 d,j). This maximum VPD was reached around 1600 h and two hours later than the maximum PAR was reached. The maximum VPD on the high VPD day was reached two hours later than the maximum on the low VPD day. The high VPD day was also characterised by both warm mornings and evenings; VPD values exceeded 2 kPa, which was higher than the day-time maximum of the low VPD day.

**Figure 2:**
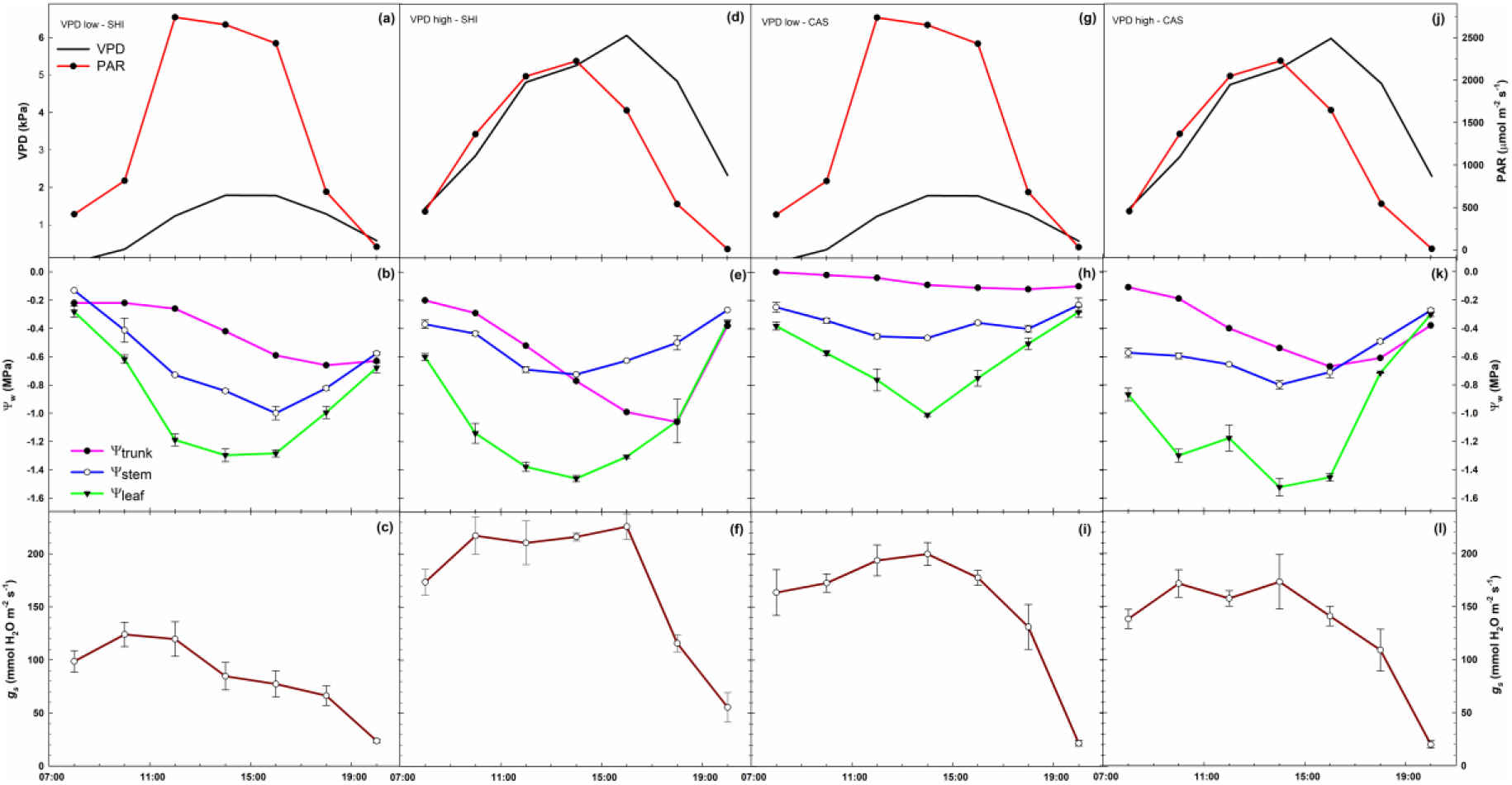
Diurnal patterns of VPD, PAR, trunk (Ψ_trunk_), stem (Ψ_stem_), leaf (Ψ_leaf_) water potentials, and leaf stomatal conductance (*g_s_*) on low and high VPD days for Shiraz (SHI) and Cabernet Sauvignon (CAS) grapevines.

Soil moisture measured using an *in situ* capacitance sensors placed approx. 30 cm below the surface and adjacent to the vine trunks indicated that the average volumetric water content (VWC) on the two measurement days were 24.0% and 30.6% on January 26 and February 17, respectively, in Shiraz; in Cabernet Sauvignon the VWC values were 28.3% and 30.3% on January 26 and February 17, respectively.

### Diurnal patterns of vine water status

Patterns of leaf (Ψ_leaf_), stem (Ψ_stem_), and trunk (Ψ_trunk_) water potentials of the same vines were quantified over the course of the two measurements days under low and high VPD conditions for both Shiraz and Cabernet Sauvignon grapevines (Fig. 2b,e,h,k). A general pattern was observed in all the vines of a gradual decrease in water potential (Ψ_w_) over the course of the day, reaching a minimum in the mid-afternoon, followed by an increase in the evening. The water potentials observed across all dates and cultivars ranged between −0.3 MPa and −1.0 MPa for Ψ_trunk_, between −0.1 MPa and −0.95 MPa for Ψ_stem_, and −0.25 and −1.5 MPa for Ψ_leaf_. The lowest Ψ_w_ values were generally observed around 1400 h each day, after which the values increased gradually.

Under low VPD (~ 1.7 kPa) conditions, Shiraz grapevines had Ψ_trunk_,Ψ_stem_, and Ψ_leaf_ daily average minimum values of −0.66, −0.96, and −1.30 MPa, respectively (Fig. 2b). Although the three Ψ_w_ values were similar in the early morning (0800 h) and late evening (2000 h), they reached maximum separation between 1200 h and 1400 h, during which time the difference between them were approx. 0.3, 0.3, and 0.6 MPa for ΔΨ_trunk-stem_, ΔΨ_stem-leaf_ and ΔΨ_trunk-leaf_, respectively. In comparison, Cabernet Sauvignon on the same day, i.e. under low VPD conditions, had Ψ_trunk_,Ψ_stem_, and Ψ_leaf_ daily average minimum values of −0.12, −0.45 and −1.01 MPa, respectively (Fig. 2h), which were higher than the values observed in Shiraz for all three metrics. Similar (but not identical) to Shiraz, the differences between Ψ_w_ values were greatest around 1400 h; difference values were approx. 0.3, 0.3 and 0.9 MPa for ΔΨ_trunk-stem_, ΔΨ_stem-leaf_ and ΔΨ_trunk-leaf_, respectively. The diurnal patterns of Ψ_w_ were somewhat different compared to Shiraz; Cabernet did not drop its Ψ_trunk_ and Ψ_stem_ as much as Shiraz under the same conditions despite its Ψ_leaf_ dropping to slightly below −1 MPa.

Under high VPD (~ 6 kPa) conditions, Shiraz grapevines dropped their Ψ_trunk_, Ψ_stem_ and Ψ_leaf_ values to −1.1, −0.7, −1.5 MPa, respectively (Fig. 2e), which were considerably lower than during the low VPD day. Under these high VPD conditions in both cultivars, patterns of Ψ_w_ early in the day were similar to the patterns observed during the low VPD day; the values of Ψ_w_ followed the expected order of Ψ_trunk_ > Ψ_stem_ > Ψ_leaf_ during the morning. However, a distinct shift in the pattern of Ψ_trunk_ was observed after approx. 1400 h: Ψ_trunk_ dropped below Ψ_stem_, reaching a minimum of −1.06 MPa around 1800 h, matching values observed for Ψ_leaf_. In comparison, Ψ_stem_ at the same time (1800 h) was −0.47 MPa. There was a noticeable recovery (increase) in Ψ_trunk_ that started soon after 1800 h, matching the values of Ψ_leaf_ during this period late in the day (1800 h – 2000 h). Under the same (high VPD) conditions, patterns of Cabernet Sauvignon Ψ_w_ were similar to that of Shiraz. Cabernet had large differences between Ψ_trunk_, Ψ_stem_ and Ψ_leaf_ early in the day, and these reached their minimum values of −0.67, −0.76, and −1.5 MPa, respectively, between 1400 – 1600 h (Fig. 2k). Much like Shiraz under similar (high VPD) conditions, there was a distinct crossing-over of Ψ_trunk_ and Ψ_stem_ around 1600 h; Ψ_trunk_ dropped to values below Ψ_stem_, remaining in this position until the end of the day. Differences between the various metrics of Ψ_w_ were more significant in Cabernet than in Shiraz; maximum difference values in Cabernet were approx. 0.1, 0.8 and 0.9 MPa for ΔΨ_trunk-stem_, ΔΨ_stem-leaf_ and ΔΨ_trunk-leaf_, respectively, while in Shiraz the corresponding differences were 0.05, 0.7, and 0.7 MPa, respectively.

Leaf stomatal conductance (*g_s_*) was measured concurrently with Ψ_w_ measurements, and on the same leaf used to measure Ψ_leaf_. Patterns of *g_s_* mirrored Ψ_w_ in both cultivars and across both measurement days under contrasting environmental conditions. In Shiraz, average *g_s_* values were highest early in the day, peaking around 124 mmol H_2_O m^−2^ s^−1^ at 1000 h under low VPD conditions, and considerably higher around 235 mmol H_2_O m^−2^ s^−1^ also at 1000 h under high VPD conditions (Fig. 2c,d). Under high VPD conditions, however, there was a precipitous decline in *g_s_* after 1600 h down to similar values observed under low VPD conditions by late day (2000 h). Within a two hour window, between 1600 h and 1800 h, Shiraz *g_s_* values dropped by an average of 49% or 110 mmol H_2_O m^−2^ s^−1^.

In Cabernet Sauvignon, *g_s_* values increased steadily from morning until around 1400 h (max. *g_s_* ~ 200 mmol H_2_O m^−2^ s^−1^) under low VPD conditions (Fig. 2i), decreasing rapidly to similar values as those observed in Shiraz under the same conditions. In contrast with Shiraz under high VPD conditions, Cabernet had lower *g_s_* early in the day until around 1400 h, after which a steady decrease was observed; *g_s_* values late in the day matched those under low VPD conditions as well as Shiraz under both environmental conditions.

### Dynamics of soil-plant-environment interactions

Diurnal patterns of environmental conditions, soil moisture, and vine water status were studied over a period of eight days between January 19 and January 26, 2021. This period encompassed a range of environmental conditions including hot and cool days (with corresponding high and low VPDs), and irrigation and precipitation events. There was an increasing temperature and VPD trend during the first three days of this period, followed by a decrease on January 22^nd^ leading to the highest VPD values observed on January 24^th^ (daily max. VPD ~ 6.7 kPa). Over this period, approx. 20 mm of irrigation was applied in the Cabernet block and 8 mm in the Shiraz block, in addition to the nearly 10 mm of precipitation received in the region on January 25^th^ (Fig. 3a). Changes in soil moisture reflected these events; VWC values in the Cabernet block increased between 10-14% with 7.6 mm of irrigation and less than 5% with the extra rainfall of 10 mm (Fig. 3b). The same rain event resulted in the soil moisture increasing by approx. 12% in the Shiraz block with the same soil type.

**Figure 3:**
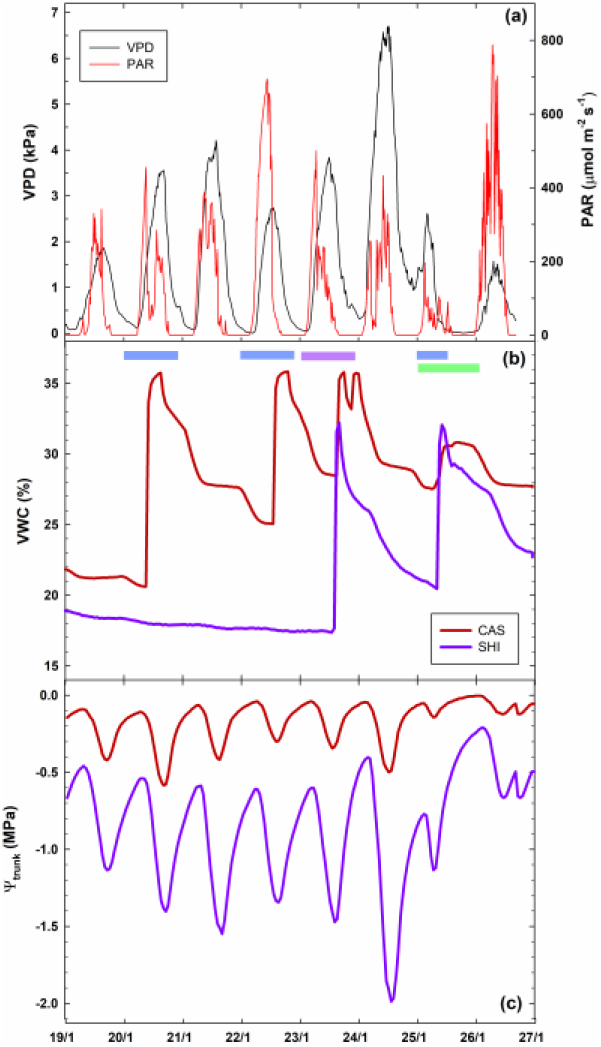
Week-long diurnal patterns of VPD and PAR **(a)**, soil volumetric water content **(b)**, and Ψ_trunk_ **(c)** for Shiraz and Cabernet Sauvignon grapevines. Blue bars on top of (b) represent irrigation events of ~7 mm for Cabernet Sauvignon, the purple bar represents irrigation of ~8 mm in Shiraz, and the green bar represents a rain event of 9.6 mm.

Trunk water potentials (Ψ_trunk_), as measured continuously by the microtensiometers, were a composite reflection of both soil moisture and environmental conditions, in particular VPD. Patterns of diurnal oscillations of Ψ_trunk_ were entrained with those of VPD albeit with time offsets on specific days (reported below; Fig. 3a,c). The daily maximum values of Ψ_trunk_ were usually observed between 0700 – 0900 h, approx. 2-3 hours after sunrise, and ranged from −0.01 to −0.10 MPa for Cabernet Sauvignon, and −0.01 and −0.6 MPa for Shiraz over the observation period (Fig. 3c). The daily minimum values of Ψ_trunk_ were usually observed between 1620 – 1820 h, and values ranged between −0.12 and −0.59 MPa for Cabernet, and −0.67 and −1.99 MPa for Shiraz.

During the first three days (of the eight-day observation period), the increasing trend of daily maximum VPD from 1.86 kPa on January 19^th^ to 4.21 kPa on January 21^st^ resulted in a concomitant decline in daily minimum Ψ_trunk_ from −0.42 to −0.59 MPa (on Jan-20) in Cabernet Sauvignon, and −1.14 to −1.55 MPa in Shiraz (on Jan-24). Soil moisture levels did not vary for Shiraz during this period, while Cabernet vines received 7.6 mm of irrigation on January 20^th^ that resulted in an increase of VWC from 22% to 36% (Fig. 3b). This irrigation event resulted in an increase of minimum Ψ_trunk_ in Cabernet on January 20-21 from −0.59 MPa to −0.43 MPa despite higher VPD levels on January 21. The Ψ_trunk_ responses of both cultivars to these conditions mirrored those of VPD; higher VPD values resulted in lower Ψ_trunk_. Shiraz Ψ_trunk_ appeared to be strongly coupled to VPD; Pearson correlation coefficients (*R*) were highly correlated with daily minimum Ψ_trunk_ and ΔΨ_trunk_ (=max. Ψ_trunk_ − min. Ψ_trunk_), whereas Cabernet was only weakly correlated for the ΔΨ_trunk_ parameter (Table 1).

**Table 1.** Pearson correlation coefficients (*R*) of daily maximum VPD vs. minimum and maximum Ψ_trunk_ values, and diurnal Ψ_trunk_ differences (Δ Ψ_trunk_ = max. Ψ_trunk_ − min. Ψ_trunk_) during the eight day observation period from January 19-26, 2021 for Shiraz and Cabernet Sauvignon. n=8.

| Parameter | Cabernet Sauvignon |  |  | Shiraz |  |  |
| --- | --- | --- | --- | --- | --- | --- |
| | Max. $\Psi_{\text{trunk}}$ | Min. $\Psi_{\text{trunk}}$ | $\Delta\Psi_{\text{trunk}}$ | Max. $\Psi_{\text{trunk}}$ | Min. $\Psi_{\text{trunk}}$ | $\Delta\Psi_{\text{trunk}}$ |
| Pearson $R$ | -0.10 | -0.60 | -0.70 | -0.35 | -0.94 | -0.96 |
| $P$-value | 0.811 | 0.115 | 0.053 | 0.393 | 0.001 | 0.000 |

In the next period of three days, two warm-to-hot days were experienced between January 23-24 with maximum VPD values of 3.84 and 6.70 kPa, respectively. On the warmest of these days, January 24, minimum Ψ_trunk_ values dropped to −0.51 and −1.99 MPa in Cabernet and Shiraz, respectively. Irrigation applied in these blocks between January 23-24 resulted in higher maximum Ψ_trunk_ values, particularly in Shiraz, increasing from −0.60 to −0.40 MPa. On the same warm day (January 24), the diurnal decline in Ψ_trunk_ in Cabernet was approx. 0.5 MPa while in Shiraz the decline was approx. 1.6 MPa (Fig. 3c). On this warm day, Ψ_trunk_ continued to drop until around 1720 h in Cabernet and until around 1820 h in Shiraz (Note: the highest VPD value was observed around 1730 h). In comparison, on cooler (low VPD) days, minimum Ψ_trunk_ values were typically observed around 1620 h and 1740 h for Cabernet and Shiraz, respectively. On these cooler days, the highest VPD is typically reached around 1600 h, close to the time of minimum Ψ_trunk_ of Cabernet, but one hour earlier than that of Shiraz, similar to the observation on the high VPD day. On cooler days with max. VPD < 2.0 kPa, diurnal changes in Ψ_trunk_ averaged 0.2 MPa for Cabernet, and 0.7 MPa for Shiraz. A marked improvement in vine water status (increase in Ψ_trunk_) was observed from January 25^th^ onward resulting from both increases in soil moisture (via irrigation and precipitation), and probably more significantly, decreases in VPD. By the end of the observation period, January 26, minimum Ψ_trunk_ values had risen to −0.12 and −0.67 MPa in Cabernet and Shiraz, respectively.

The relationships between VPD and Ψ_trunk_ for Cabernet and Shiraz for the period December 14, 2020 to February 10, 2021 (0600-2000 h) are presented in Fig. 4. There was a significant difference between cultivars in the sensitivity of vine water status to VPD (as indicated by the slopes of the regression lines) during the two-month peak summer period. Shiraz was more sensitive than Cabernet to changes in atmospheric conditions, dropping its Ψ_trunk_ by approx. 0.17 MPa kPa^−1^. In comparison, Cabernet reduced its Ψ_trunk_ by approx. 0.07 MPa kPa^−1^.

**Figure 4:**
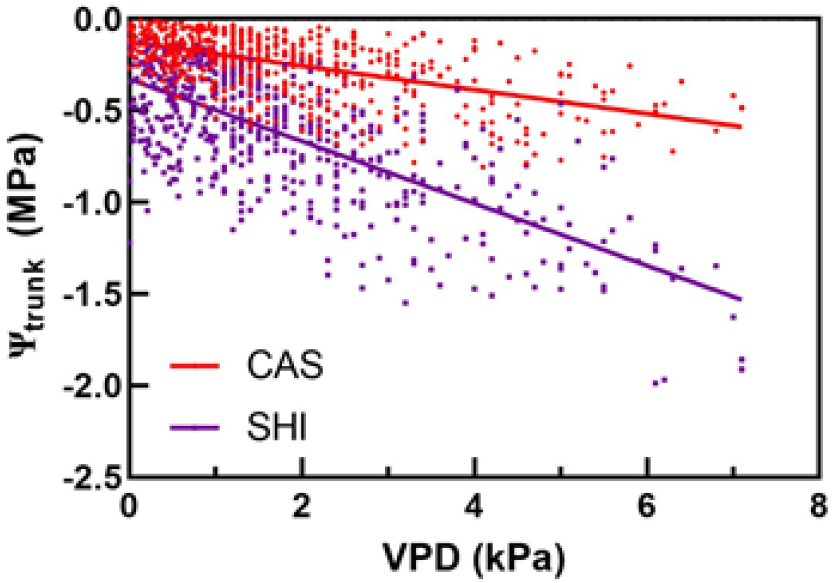
Linear regression analysis of VPD vs Ψ_trunk_ of Shiraz and Cabernet Sauvignon grapevines over the measurement period. Regression equations: Shiraz: Ψ_trunk_ = −0.3272 - 0.1696*VPD, R^2^ = 0.51; Cabernet Sauvignon: Ψ_trunk_ = −0.1247 - 0.065*VPD, R^2^ = 0.33. *P*-value for differences between slopes: < 0.001.

Time-lagged cross correlation analysis (TLCC) was used to analyse the time series datasets of VPD and Ψ_trunk_ across two days with contrasting environmental conditions or VPDs: January 26, 2021 (low VPD day; daily max. VPD ~ 1.6 kPa) and February 17, 2021 (high VPD day; daily max. VPD ~ 6.7 kPa). On the low VPD day, the diurnal pattern of VPD followed the typical trend increasing from around 0900 h to its maximum around 1500 h after which a gradual decline commenced until reaching its minimum around 2000 h (Fig.5a).

Ψ_trunk_ followed a similar but mirrored trend, increasing in the early hours of the day, then decreasing from around noon to its minimum value of around 1800 h before rising again (Fig. 5b). Cabernet Sauvignon (CAS) and Shiraz appeared to have very similar diurnal patterns of Ψ_trunk_ despite differences in absolute Ψ_trunk_ values. TLCC analysis between VPD and Ψ_trunk_ revealed that CAS had the minimum Pearson Product Moment (normalised cross correlation) coefficient (XCC) at an offset of −9 units or −180 min (each offset unit = 20 min), indicating that Ψ_trunk_ lagged VPD by 180 min (Fig. 5c). Similarly, Shiraz Ψ_trunk_ lagged VPD by 220 min (Fig. 5d).

**Figure 5:**
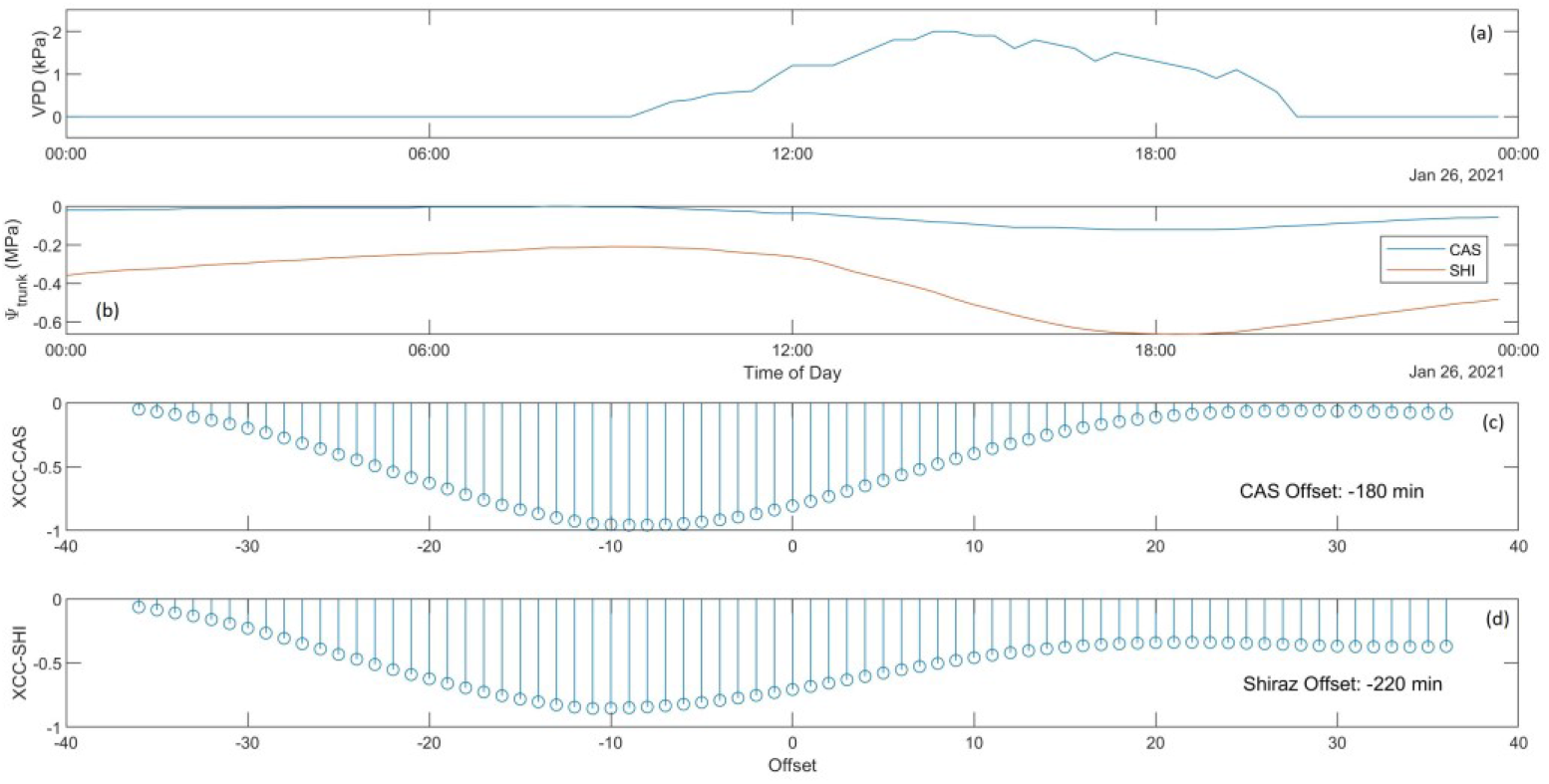
Diurnal plot on a low VPD day (maximum VPD ~ 1.6 kPa) of **(a)** VPD, **(b)** Ψ_trunk_ for Shiraz and Cabernet Sauvignon, and Pearson Product Moment (cross correlation) coefficient values for Cabernet **(c)** and Shiraz **(d)** showing leading or lagging of Ψ_trunk_ in response to changing VPD.

On the high VPD day of January 24, the maximum VPD of the day of approx. 6.7 kPa was reached around 1700 h (Fig. 6a), while the minimum Ψ_trunk_ in both Shiraz and CAS were much lower than those on the low VPD day, reached around 1800 h (Fig. 6b). TLCC analysis revealed that CAS Ψ_trunk_ lagged VPD by 80 min (Fig. 6c) while Shiraz Ψ_trunk_ lagged VPD by 120 min (Fig. 6d), both lower than the time lags observed on the low VPD day.

**Figure 6:**
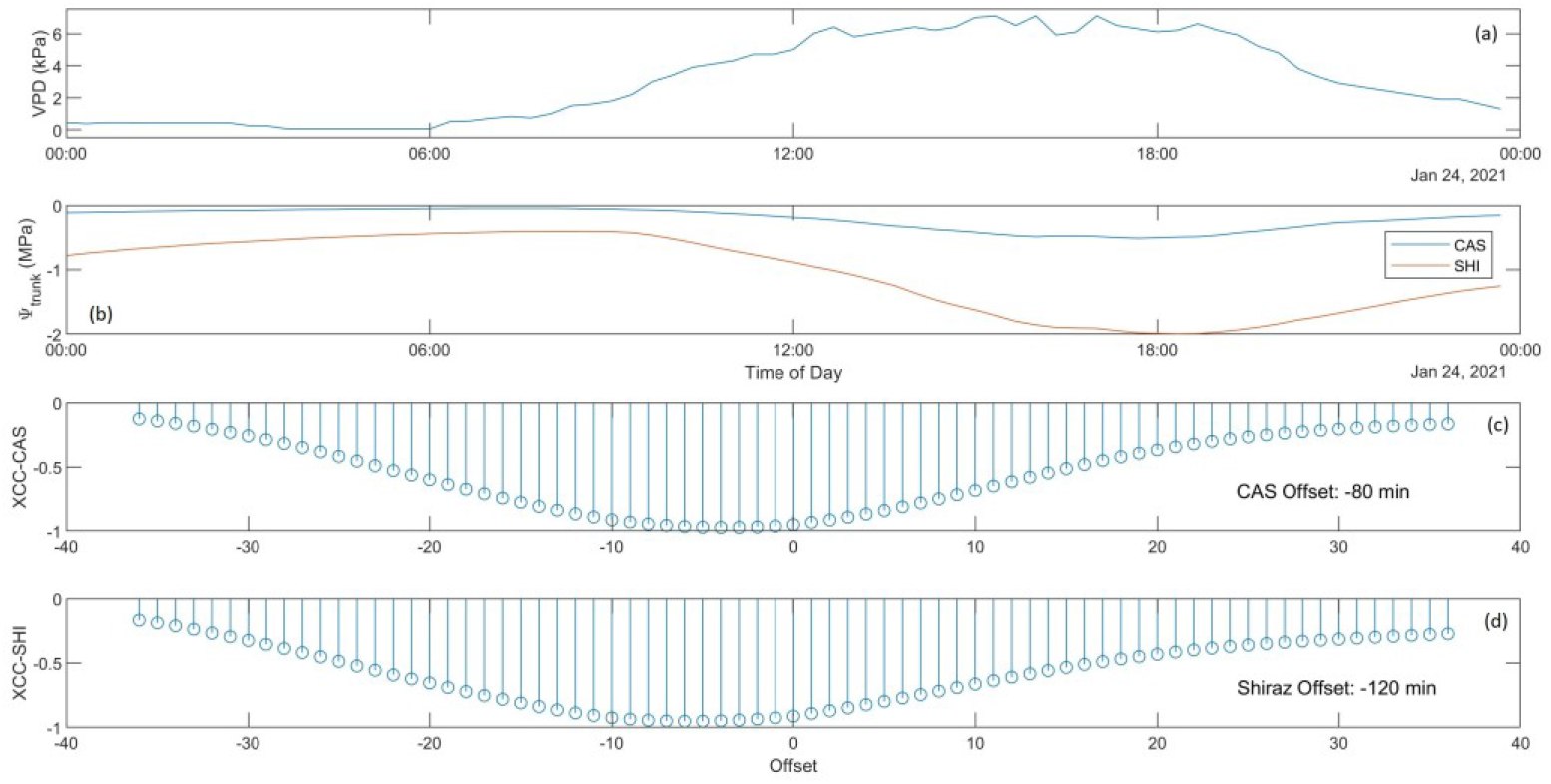
Diurnal plot on a high VPD day (maximum VPD ~ 6.7 kPa) of **(a)** VPD, **(b)**Ψ_trunk_ for Shiraz and Cabernet Sauvignon, and Pearson Product Moment (cross correlation) coefficient values for Cabernet **(c)** and Shiraz **(d)** showing leading or lagging of Ψ_trunk_ in response to changing VPD.

## Discussion

Microtensiometers (MT) provide a rapid and continuous *in situ* measurement of water potential in virtually any matrix, and in our context, in woody grapevine trunks. *In situ* measurements of plant water potential provide valuable estimates of their water status, which are critical for determining irrigation schedules, i.e. when and how much water to apply, for irrigated crops. Although a number of other plant-based sensors currently exist, only very few measure water potential continuously, and those can be challenging to install and unsuitable for long term measurements in the field. MTs overcome some of these limitations due to their small form factor thereby allowing them to be embedded within the woody structures of the plant. This location minimises the risk of damage to the sensing elements from farm machinery, as well as environmental and biotic factors. This first report on the field testing of MTs in mature grapevine trunks under varying environmental conditions provides valuable data of Ψ_trunk_ on their performance in comparison to traditional methods of water potential measurement, e.g. leaf pressure chamber. To the best of our knowledge, this is the first report of field testing results of MTs. To date, the only other reports of continuous *in situ* water potentials are reported using stem psychrometers (Dixon and Tyree, 1984; McBurney and Costigan, 1984; Tran *et al.*, 2014) and an osmometric sensor (Meron *et al.*, 2015).

### Diurnal patterns of vine water status

Plant water potential reflects the dynamic interplay between incident solar radiation, VPD, and soil moisture availability, as well as several internal factors within the plant that regulate water transport from the roots to leaves. In the present study, diurnal measurements of leaf, stem and, for the first time, trunk water potentials were conducted on several days characterised by contrasting environmental conditions with low (< 2.0 kPa) and high (> 6 kPa) VPDs (Fig. 2). During this time, soil moisture content (volumetric water content), as measured by a capacitance sensor, varied two-fold from approx. 18% to 36% in a well-drained soil. Previous studies documenting diurnal patterns of leaf water potential (Ψ_leaf_) showed similar trends to the present study; a hyperbolic or U-shaped curve is typically observed with the highest values in the early hours of the day and lowest values in the afternoon in grapevine (Smart, 1974; van Zyl, 1987) and other woody horticultural crops (Klepper, 1968; Olsson and Milthorpe, 1983; Smart and Barrs, 1973). In *Vitis vinifera* cv. Colombar (grafted onto 99R rootstock), sunlit leaves had lower Ψ_leaf_ compared to shaded leaves, and the former reached the lowest Ψ_leaf_ earlier in the day (1200 h for sunlit compared to 1400 h for shaded), but only when there was a deficit in soil moisture (van Zyl, 1987). In California, Williams and Baeza (2007) observed in several red grape cultivars that the lowest Ψ_leaf_ was reached around 1600 h when the vines were either well-watered, receiving irrigation equal or higher than full replacement (100%) of crop evapotranspiration (ET_c_), or deficit irrigated at 40% ET_c_. In comparison, Thompson Seedless reached its lowest Ψ_leaf_ by around 1300 h when well-watered and by 1000 h under non-irrigated conditions. The Williams and Baeza study unfortunately did not report on the early morning or late day recovery patterns in Ψ_leaf_ of these vines so difficult to compare with the current study. In Tempranillo grapevines receiving 70% ET_c_, Ψ_leaf_ and Ψ_stem_ were observed to be at their lowest values around 1400 h and at a similar time to the highest VPD of the day (Cole and Pagay, 2015), corroborating with observations in the present study.

Freeman *et al.* (1982) found in Chardonnay and Carignan that the lowest Ψ_leaf_ was reached between 1500 h – 1600 h in Davis, California conditions. In the Chasselas cultivar in Switzerland, Zufferey and co-workers found that the diurnal patterns of Ψ_stem_ showed a similar U-shaped pattern, and reached a minimum value around 1430 h on a high VPD day (~ 3.0 kPa) independent of soil moisture and at the same time on a low VPD day (~ 1.5 kPa) in irrigated vines (Zufferey *et al.*, 2011). However, on a low VPD day without irrigation, the vines reached their minimum Ψ_stem_ values only around 1600 h likely due to the low transpiration rates. The effect of soil moisture on diurnal Ψ_leaf_ patterns is quite marked; irrigation of Shiraz in a warm climate resulted in the highest transpiration rates and lowest Ψ_leaf_ around 1200 h and 1500 h, respectively (Smart, 1974). In comparison, water-stressed vines reached their lowest Ψ_leaf_ also around 1500 h, but the highest transpiration rates were reached around 1100 h. Our diurnal observations indicated that the lowest Ψ_leaf_, Ψ_stem_, and Ψ_trunk_ were reached around 1400 h, 1600 h and 1800 h, respectively, on low VPD days, whereas these times were advanced on the high VPD day, particularly for Ψ_stem_ and Ψ_trunk_. Williams and Baeza (2007) suggested that the influence of VPD on Ψ_leaf_ decreases as soil moisture decreases.

Modeling of diurnal patterns of Ψ_leaf_ predicted that the lowest values are likely to be reached around 1400 h, approx. two hours after the highest leaf transpiration rates are reached (Katerji et al. 1986). This transient (or delay) in Ψ_stem_ response can be attributed to the contributions of both plant tissue water (~ 13%) and root uptake of soil moisture (~ 87%) to the transpiration stream. These modelled patterns compare favourably with those obtained from lysimeters; maximum transpiration rates were reached between 1200 h and 1400 h (Williams *et al.*, 2012). The same study found that higher vine sizes or crop factors result in not only increased transpiration rates, as expected, but also a delayed peak by as much as two hours. Similar patterns of vine transpiration were observed in studies using sap flow and thermal dissipation sensors (Braun and Schmid, 1999; Pagay and Skinner, 2018). The diurnal measurements of leaf stomatal conductance (*g_s_*) in the present study indicated that the highest values were reached in the afternoon on low VPD days and late morning on high VPD days. Reductions in *g_s_* following this peak resulted in increases in Ψ_leaf_ and Ψ_stem_, but not Ψ_trunk_. This lack of response from Ψ_trunk_ might indicate a level of buffering of water potential in the woody organs of the plant, in this case the trunk, where xylem vessels are surrounded by parenchymal cells that can contribute water to the transpiration stream, the so-called ‘capacitance effect’ (Salomón *et al.*, 2017; Waring and Running, 1978).

Environmental conditions played a key role in determining patterns and values of Ψ_w_; both Ψ_leaf_ and Ψ_stem_ were strongly influenced by VPD. Consistent with other studies, our study found a strong negative relationship between Ψ_trunk_ and VPD (Fig. 4). In California, across three cultivars, Williams and Baeza (2007) found that the slope of the VPD-Ψ_leaf_ relationship was 0.079 MPa kPa^−1^while for the VPD-Ψ_stem_ relationship was 0.068 MPa kPa^−1^. In comparison, our study found slopes for VPD-Ψ_trunk_ in Cabernet Sauvignon to be 0.07 MPa kPa^−1^ while for Shiraz this slope was higher at 0.17 MPa kPa^−1^. The higher sensitivity of Shiraz grapevines to VPD might reflect its relatively anisohydric behaviour that has been previously reported (Dayer *et al.*, 2020; Schultz, 2003), with a weaker regulation of stomatal conductance (*g_s_*) under either declining soil moisture or increasing VPD. In contrast, Cabernet Sauvignon’s lower Ψ_trunk_ sensitivity to VPD suggests a tighter coupling to the environment – both soil moisture and VPD.

### Temporal coupling of plant water status and atmosphere

Continuous measurements of plant water status provide a valuable dataset for the analysis of its temporal patterns as well as relationships with the environment, both soil and plant. Our characterisation of the dynamic nature of Ψ_trunk_ in Shiraz and Cabernet Sauvignon grapevines using MTs was done both inter- and intra-day (diurnally). Furthermore, using a time-lagged cross correlation statistical approach, we conducted a detailed temporal analysis of the interrelationships between VPD and Ψ_trunk_ that revealed the differences between the two grapevine cultivars in their response to different environmental conditions. Cross-correlation analysis has been used for signal processing and pattern recognition in diverse fields including spectroscopy, seismology, finance, and quantum information processing (Podobnik and Stanley, 2008). In plants, the response of water potential to the environment has been characterised previously, but comparisons of different cultivars using *in situ* measurements of Ψ_w_ have not been reported hitherto. The slower transient or response of Ψ_trunk_ to changes in VPD over the course of a day in Shiraz compared to Cabernet (Fig. 5) may reflect its greater capacity for root water uptake and/or increased capacitance from the parenchymal cells surrounding xylem vessels. Although not specifically measured in this study, it is likely that the Shiraz vines had bigger root systems (biomass) than Cabernet based on the visually larger canopies and higher leaf area index (LAI) values (Shiraz LAI ~ 7.4; Cabernet LAI ~ 5.6; average LAI data obtained from an accompanying study on 32 vines per cultivar).

Under environmentally demanding (high VPD) conditions, the Ψ_trunk_ of both cultivars were more strongly coupled to VPD with shorter transients. The more rapid vine response under high VPD conditions may result from higher transpiration rates and more open stomata, which allowed the Ψ_trunk_ in Shiraz to drop to nearly −2 MPa compared to −0.6 MPa on the low VPD day. In comparison, the relatively modest reduction of Cabernet Ψ_trunk_ of ~ 0.4 MPa when the VPD increased may indicate near-isohydric behaviour. This observation is consistent with other reports of the stomatal behaviour of Cabernet Sauvignon (Collins and Loveys, 2010), but in contrast to other reports that Cabernet Sauvignon has anisohydric behaviour (Suter *et al.*, 2019).

The stem potential sensor based on osmometry tested in tangerine trees in Israel provided diurnal Ψ_stem_ values that correlated temporally with trunk temperature and evapotranspiration (ET), showing a stem water potential lag of approx. five hours with both parameters (Meron *et al.*, 2015). The same sensor used in peach trees showed a lag of five hours with destructive, pressure chamber measurements of Ψ_stem_ and a similar (ca. five hour) lag from ET. Our results indicate that the time lag of Ψ_trunk_ was cultivar- and VPD-dependent; in the order of 3-4 hours on a low VPD day and 1-2 hours on a high VPD day with Cabernet at the low end of these ranges.

### Measurement of predawn water potential and its timing

The predawn water potential (Ψ_pd_) is considered a reliable indicator of root xylem pressure potential (Ψ_p_) or soil matric potential (Ψ_m_) based on the equilibration of Ψ_w_ between the plant and soil when the canopy transpiration rate (*E*) is zero or negligible (Ameglio *et al.*, 1999; Jones, 2007). The Ψ_pd_ is influenced by soil water content and distribution, root area/distribution, and soil and root hydraulic conductivities (Garcia-Tejera *et al.*, 2021). It should be noted that Ψ_pd_ is not a reflection of the average soil Ψ_w_ across the rhizosphere, but rather the Ψ_w_ of its wettest portion (Pagay *et al.*, 2016; Schmidhalter, 1997). Furthermore, Ameglio *et al.* (1999) reported that Ψ_pd_ may not reflect plant Ψ_w_ and hence has limited application for irrigation scheduling of crops. On a diurnal basis, our observation that the maximum Ψ_w_ in the plant (based on Ψ_trunk_), widely accepted as the Ψ_pd_ (Boyer, 1995), occurred around 0700-0800 h, 2-3 hours after sunrise and that reported in the literature (Chone *et al.*, 2001; Correia *et al.*, 1995; Williams and Araujo, 2002). The fundamental requirement for the equilibration of plant and soil Ψ_w_ is that (nocturnal) transpiration is zero or near zero, which is sometimes not the case in well-watered crops in warm-to-hot climates (Pagay, 2016). Previous studies characterising diurnal canopy *E* showed that *E* reaches its minimum value (zero or near-zero) around 0100 h under low VPD conditions and around 0600 h under high VPD conditions, under well-watered conditions (Pagay, 2016). Using sap flow sensors in grapevines, Braun and Schmid (1999) observed that the minimum canopy *E* for the day was reached around 2100 h, and *E* did not start increasing until approx. 0900 h, several hours after dawn, the next day. Differences between the times of minimum *E* and highest Ψ_w_ diurnally may be the result of hydraulic resistances in the soil-plant continuum, which are influenced by soil type, soil moisture, ambient VPD, and root biomass and distribution (Garcia-Tejera *et al.*, 2021; Sato *et al.*, 2006). Our observations of maximum plant Ψ_w_ being reached in the early morning, between 0700-0800 h, are consistent with another study reporting that Ψ_stem_ does not start decreasing from its maximum diurnal value until the early morning, approx. 0700 h (Cole and Pagay, 2015), but in contrast to another report that Ψ_pd_ decreases from 0330 h (Carbonneau *et al.*, 2004). The observation has implications for the timing of measurement of Ψ_pd_, if used as a metric for crop irrigation scheduling as recommended previously (Stricevic and Caki, 1997). Donovan *et al.* (2001) found that Ψ_pd_ and Ψ_p_ in several woody plants may not reflect Ψ_m_ even under well-watered conditions without nocturnal transpiration. This was hypothesized to be due to the accumulation of high concentrations of solutes in the leaves, although grapevines have only modest levels of osmotic adjustment compared to many other woody horticultural crops (Rodrigues *et al.*, 1993).

### Which plant water potential metric to use for irrigation scheduling?

MTs offer yet another plant water status metric, Ψ_trunk_, that has been shown in this study to have a different range of values compared to conventional measures of Ψ_leaf_ and Ψ_stem_. These conventional metrics have been well characterised for irrigation scheduling and thresholds have been suggested in the literature (Deloire and Heyns, 2011; Romero *et al.*, 2010).

#### i. Leaf water potential

The Ψ_leaf_ is a convenient measurement with the use of a leaf pressure chamber, although time and labour intensive. The inherent variability of leaves in a plant (and even within a shoot or branch) make this metric highly variable in its value (McCutchan and Shackel, 1992). Jones (1990) suggested that Ψ_leaf_ may be an erroneous indicator of plant water status as Ψ_leaf_ homeostasis may occur under different soil moisture and environmental conditions. This homeostasis in Ψ_leaf_ is exemplified no better than in plants that lie at the opposite ends of the isohydric-anisohydric spectrum. Tardieu and Simonneau (1998) found that in sunflower and barley, characterised as anisohydric where there is weak coupling between soil moisture and stomatal conductance (*g_s_*), Ψ_leaf_ declined in relation to VPD and soil moisture. In contrast, maize, characterised as near-isohydric, where stomatal closure occurs with declining soil moisture, Ψ_leaf_ was virtually unchanged under declining soil moisture until near death. Grapevines cultivars also have been shown to vary in their homeostasis in Ψ_leaf_ under declining soil moisture. For example, Syrah (syn. Shiraz) was shown to be relatively anisohydric compared to Grenache, which was near-isohydric (Schultz, 2003). The differential response between cultivars is thought to be related to leaf and xylem abscisic acid and hydraulic regulation (Dayer *et al.*, 2020). Based on these physiological responses, the use of Ψ_leaf_ for irrigation scheduling of relatively isohydric plants may underestimate their true water stress and therefore irrigation requirements potentially leading to a vicious cycle.

#### ii. Stem water potential

The Ψ_stem_ overcomes some of the liabilities associated with Ψ_leaf_, particularly that of leaf-level variability with a shoot or branch, as it integrates all the leaves of that shoot/branch, and is therefore less variable than Ψ_leaf_ (Williams and Araujo, 2002). The Ψ_stem_ was also shown to be a more sensitive indicator of plant water status than Ψ_leaf_ (Garnier and Berger, 1985) and could reliably discriminate soil moisture deficits (Chone *et al.*, 2001) earlier than both Ψ_leaf_ and Ψ_pd_ (Selles and Berger, 1990). Despite these advantages, from a practical standpoint, Ψ_stem_ is somewhat more involved, requiring leaves to be enclosed in opaque bags to stop transpiration and allow Ψ_leaf_ and Ψ_stem_ to equilibrate; this needs to be done at least one hour prior to measurement in the pressure chamber (Chone *et al.*, 2001). This delayed measurement is not amenable to rapid measurements or automation, which applies to Ψ_leaf_ also.

#### iii. Trunk water potential

The Ψ_trunk_ is arguably the most stable of these three metrics, integrating all the leaves of the plant in a stable tissue that is relatively unaffected by external factors as are Ψ_leaf_ and Ψ_stem_. Our measurements of these three vine Ψ_w_ metrics indicated that, in some instances, Ψ_trunk_ tended to be nearly 1 MPa higher than Ψ_leaf_, indicative of the high hydraulic resistances between the trunk and leaves. Previous reports have shown that the highest hydraulic resistance in this pathway lies in the leaf, representing as much as 30% of the overall resistance in the plant (Sack *et al.*, 2003), likely due to the fewer and narrower xylem vessels in this section of the pathway compared to distal sections. The Ψ_trunk_ was also susceptible to the least fluctuations diurnally, although this was only shown to be true under low VPD conditions (Fig. 2). Its central location in the plant between the roots and leaves, as well as buffering of xylem water status (pressure potential) via capacitance from adjoining parenchymal cells and secondary xylem (Meinzer *et al.*, 2009) would be plausible reasons for the stability of the trunk’s water status. However, we observed that Ψ_trunk_ responded to changes in soil moisture (via irrigations) less rapidly than to changes in VPD (Fig. 3), which suggests that the trunk may be well-coupled to the leaves despite the high hydraulic resistances in the leaf petioles. A related and somewhat surprising observation was made under high VPD conditions: we consistently observed the crossing over of the Ψ_trunk_ and Ψ_stem_ lines in the mid-afternoon (Fig. 2e,k). The lower Ψ_trunk_ value (compared to Ψ_stem_) during warm afternoons indicates that the trunks of both cultivars were under considerably more water stressed than the stems and similar to the leaves late in the afternoon. A plausible explanation for this response is that the roots may be under water stress owing to transient water deficits at the soil-root interface due to high transpiration rates under high VPD conditions. Pagay *et al.* (2016) reported that, under high VPD conditions, low plant water potentials could result, if the capillary conductivity of soils in the rhizosphere is inadequate to support high canopy transpiration rates.

Continuous measurements of Ψ_trunk_ using *in situ* microtensiometers, which were demonstrated in field-grown plants for the first time in this study, offers a convenient measurement of plant water status for irrigation scheduling. Furthermore, these *in situ* measurements of plant water potential provide a powerful tool for physiological studies of plant hydraulics in a dynamic environment, for example, studies on the limiting water potentials of plants as well as those involving cavitation and embolism recovery dynamics. Microtensiometers are also amenable to automation, for example to automate irrigation scheduling via a decision support system in which thresholds of Ψ_trunk_ are pre-programmed in irrigation controllers for various crop phenological stages and that also incorporate other relevant environmental parameters such as weather forecast and soil moisture data for precision irrigation. The use of published Ψ_stem_ or Ψ_leaf_ thresholds to drive irrigation decisions should be based on measurements of the specific metric for which the threshold has been developed; a translation of those values to Ψ_trunk_ would not be appropriate due to physiological, hydraulic and anatomical differences between plants.

## Data availability statement

The data supporting the findings of this study are available upon request from the author.

## Acknowledgements

The author thanks the following individuals for assistance with the project: Dr Franziska Doerflinger (Plant and Food Research Australia), Dr Michael Santiago (FloraPulse), Mr Popolopoulos, Felipe Canela, and Rochelle Schlank. The project was supported by funding from Wine Australia (project: UA 1803-1.3), and in-kind support by Katnook Estate and Wynns Coonawarra Estate.

